# Boosting Bug Farms: A Meta-Analysis on Probiotic Effects in Insect Rearing

**DOI:** 10.1101/2025.04.04.647199

**Authors:** Julia Häbermann, Bernardo Antunes, Hajo Haase, Jens Rolff, Claudia Keil, Charlotte Rafaluk

## Abstract

Interest in insects and food as feed is rapidly growing. With this, however, comes a move towards mass rearing and industrial scale production. This upscaling and industrialisation of the rearing process is likely to result in high density rearing, which in itself facilitates disease spread (Tarwater and Martin, 2001). Furthermore, the fact that these insects are likely to be genetically closely related further increase the risk of infectious disease outbreak (Ekroth et al., 2019; Gibson and Nguyen, 2021). In tackling this risk, it is important that we do not resort to the mass use of antibiotics that has been seen in the livestock industry. Instead, alternative rearing practices should be developed. A practice that has received a huge increase in attention in the last years is the idea that supplementation of insect feed with probiotics could improve insect health and prevent pathogen spread. We carried out a meta-analysis to systematically analyse the data on probiotic supplementation of insects reared for food and feed available to date. The most commonly measured response to probiotic supplementation was body weight gain followed by microbiome diversity. For body weight gain, we collected 71 effect sizes from 28 studies and for microbiome diversity ten effect sizes from six studies. We found that overall, probiotics tended to increase insect growth rate but did not significantly impact microbiome diversity. Our analysis also highlighted two key literature gaps. To date, data are only available on one of the four insect species able to be sold as food in the EU, *Tenebrio molitor*. Furthermore, there are currently only very few studies that have looked at protection against pathogens by probiotic bacteria in insects reared for food and feed. So far, the data look promising, but data on more insect species and looking at inhibitory effects against pathogens are urgently needed.

## Introduction

Insect protein is rapidly gaining attention as a future food source (Hazarika and Kalita, 2023; van Huis and Gasco, 2023). This is largely because insects as a protein source represent a climate friendly and sustainable alternative to traditional meat. Around 34% of global greenhouse gas production comes from the food industry (Crippa et al., 2021), 72-78% of which come from meat production (Springmann et al., 2018). Insect rearing results in the emission of a fraction of the greenhouse gasses produced during traditional meat production. For example, the production of 1kg cricket protein produces 99% less greenhouse gas than 1kg of beef protein and overall insect rearing results in just 6% of the greenhouse gas emissions per kg body weight released during cattle rearing for beef (Jafir et al., 2024). Under the EU novel food legislation (EU) 2015/2283, four insect species, *Locusta migratoria, Acheta domesticus, Alphitobius diaperinus* and *Tenebrio molitor,* can be reared for and sold as human food in the EU (Precup et al., 2022). This rapidly growing market presents huge benefits for the equitable and sustainable production of food for a global human population that is expected to exceed 10 million by 2025 (United Nations Department of Economic and Social Affairs, Population Division, 2022). It also brings with it potential challenges (Maciel-Vergara et al., 2021), notably, an increased risk of disease spread.

Rearing of insects on the scales necessary to feed the growing human population requires mass rearing at high densities. High density cultures of genetically similar animals are known to present a particularly high risk for disease spread within populations (Ekroth et al., 2019; Gibson and Nguyen, 2021). Furthermore, insects can be reared on a range of organic side streams (Broeckx et al., 2021; van Huis, 2013; Van Peer et al., 2021; Vrontaki et al., 2024), which while reducing the environmental impact further, introduces a diversity of bacteria into the digestive tract of the insects which are likely to range from the potentially beneficial to the potentially harmful (Marzoli et al., 2024; Savio et al., 2024a; Wynants et al., 2019). The threat of potential pathogens in cultures of insects being reared for food and feed is twofold. There are pathogens that threaten the insects themselves, including entomopathogenic fungi (Dahal et al., 2022), bacterial entomopathogens, such as *Bacillus thuringiensis* (Savio et al., 2024b) and viruses (Duffield et al., 2021). Human food pathogens, in particular *Salmonella enterica* and *Bacillus cereus* have also been found in the guts of edible insect species (Fasolato et al., 2018; Marzoli et al., 2024; Wynants et al., 2019). As insects, unlike traditional livestock are generally prepared for human consumption with their guts intact, the presence of these bacteria in the insect digestive tract poses a potential threat of transmission to the human consumer.

Despite the threat posed by potential bacterial and fungal pathogens to insect cultures, it is imperative that mass prophylactic administration of antimicrobials does not become standard practice. The widespread use of antibiotics in the meat farming industry (Ghimpe_eanu et al., 2022) has significantly contributed to the global risk presented by antimicrobial resistance (AMR) (Djordjevic et al., 2024). The insect microbiota is known to be a potential source of AMR genes (Raka et al., 2024; Rawat et al., 2023). Such genes are currently found in lower abundance in insects than livestock (Raka et al., 2024) but would likely rapidly increase in frequency if selection pressure through the prophylactic use of antibiotics was applied. It is critical that the mistakes made in livestock farming are not repeated in mass insect rearing and that alternative methodologies are developed to protect insect cultures and reduce the risk of spreading antibiotic resistant bacteria into the food and feed system.

Recently, probiotics have been receiving increasing attention as a potential strategy to increase the overall health of insect cultures (Dahal et al., 2022; Grau et al., 2017; Savio et al., 2022). We know from research in the field of evolutionary ecology that bacteria with potentially protective effects, also known as “defensive microbes”, have the potential to provide stable, long-term defence against pathogens (Armitage et al., 2022; King and Bonsall, 2017; Vorburger and Perlman, 2018). For example, the gut microbe *Enterococcus mundtii* works together with its host *Galleria mellonella* to control the host microbiome during meta-morphosis, protecting against the proliferation of pathogenic bacteria (Johnston and Rolff, 2015). Similarly, the bacterial symbiont, *Hamiltonella defensa*, protects its aphid host against parasitoid wasp attack (Kaech et al., 2022; Kwiatkowski et al., 2012; Wu et al., 2022), with implications for pest control (Donner et al., 2023).

Many probiotics fall into the category of “defensive microbe”, in that they provide protection against infection (Corr et al., 2007; Deriu et al., 2013; Do et al., 2024; Fukuda et al., 2011; Piewngam et al., 2021, 2018). These probiotics can be harnessed not just to support insect growth and nutritional health, but to protect against infection and disease spread (Ford and King, 2016). In contrast to antibiotics, probiotics or defensive bacteria present a dynamic and (co)evolving defensive agent, that can change to counter adapt to a pathogen evolving resistance (Ford et al., 2017, 2016; Kwiatkowski et al., 2012). Furthermore, many human probiotics have been shown to have beneficial effects on edible insects (e.g. (Lecocq et al., 2021; Milanović et al., 2021)), meaning that supplementation may not only benefit the health of the insect cultures but has the potential to provide benefits to the human consumer as well.

Over the last three to four years, there has been a burst of publications on probiotic supplementation in edible insect species. Here we use a meta-analytic approach to summarise and systematically review these data, draw conclusions on what we know so far and highlight literature gaps where we find that more research is needed.

## Materials and Methods

### Literature search

Our initial aim was to find all papers reporting the results of studies where edible insects had been supplemented with probiotics. We used the search terms including “*Tenebrio molitor* probiotics”, “*Bombyx mori* probiotics”, “*Hermetia illucens* probiotics”, “*Apis mellifera* probiotics”, “*Acheta domesticus* probiotics”, “*Alphitobius diaperinus* probiotics”, “*Locusta migratoria* probiotics” and “edible insect probiotics” to search the databases: PubMed, Google Scholar and Web of science. We then looked at the papers citing the papers we had found and checked the references lists of all discovered papers for more potential studies.

We noted that the most commonly measured parameters in the studies found were insect growth rate and microbiome diversity (often measured as the Shannon index) and therefore decided to carry out two meta-analysis the first asking whether probiotic supplementation influenced insect growth rate and the second asking whether probiotic supplementations influenced insect microbiome diversity.

### Inclusion criteria

Papers meeting the following criteria were included in the meta-analyses:

- The paper was published in a peer reviewed journal;
- The paper was written in English;
- The host species was insect species either already registered as a novel food within the EU, which is under evaluation as a novel food in the EU or is regularly consumed as protein source outside of the EU;
- The paper presented data on, for meta-analysis one, insect growth rate or, for meta-analysis two, microbiome diversity, for both a probiotic treated and a control group;
- Means and standard errors could be extracted, calculated or estimated from the presented data either from graphs, raw data or other measures of average and spread of the data.

### Statistical analysis

All analyses were carried out in R (v4.4.1; R Core Team). For both the growth rate and diversity data sets, we calculated Hedge’s g effect sizes and confidence intervals using the “esc” package in R (Lüdecke, 2019). We then carried out a multivariant meta-analyses using the rma.mv function in the “metafor” package (Viechtbauer, 2025) with “study”, “insect” and “probiotic species” as random factors except where insect and probiotic species were explicitly being tested as moderator variables. For each of the two data sets we carried out a four moderator variable analyses which each of the follow moderators as a fixed factor. We tested whether 1) insect species, 2) probiotic species, 3) whether the probiotic was a lactobacillus or not and 4) whether the probiotic was gram positive, or gram negative had an impact on the magnitude or direction of the effect. We plotted funnel plots to visually assess publication bias and calculated fail safe N values to estimate the number of datasets that would need to be added to the analysis to change the outcome (Orwin, 1983).

## Results

### Growth rate

#### Overall model for growth rate

Through our literature search we collected a total of 71 datasets from 24 publications presenting data on growth rate under control and probiotic supplemented conditions and that met our inclusion criteria (Table 1). These data showed that, overall, supplementation with probiotics had a positive effect on growth (multi-variant meta-analysis: estimate = 2.7036 (ci.lower = 1.2541/ ci.upper = 4.1532), z = 3.6556, p = 0.0003), suggesting that overall supplementation with probiotics leads to heavier insects (Figure 1). We noticed during our data collection that some feeding material was fermented, this was specifically the case for black soldier fly larvae. As fermentation potentially results in probiotic enrichment, we noted these cases in Table 1.

**Figure 1.**
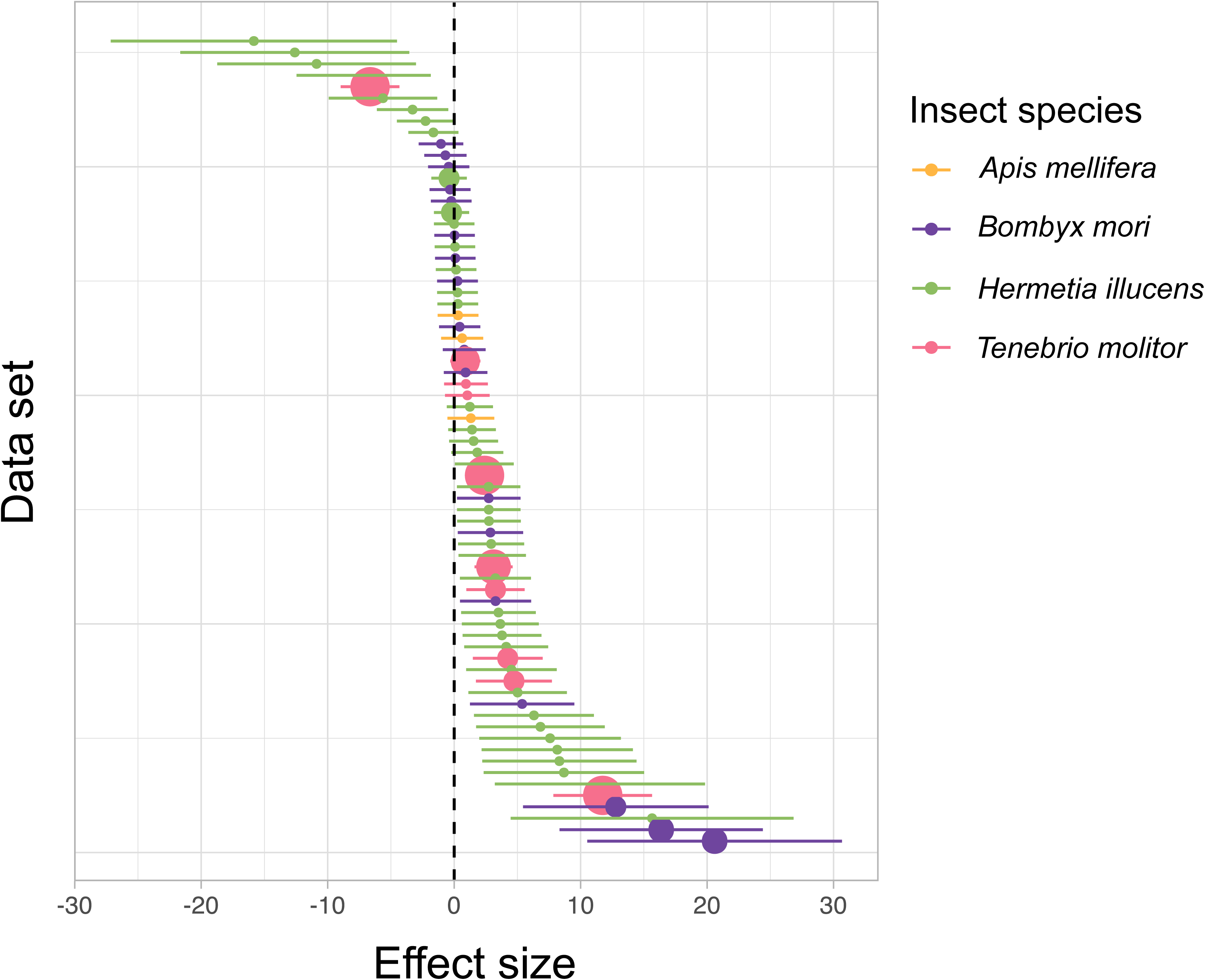
Forest plots showing effect sizes and confidence intervals for differences in insect growth with and without probiotic supplementation with effect sizes and 95% confidence intervals for each data set included in the study, negative effect sizes show a lower body weight in insects reared with probiotics than reared without and positive effect sizes show increased growth in insects reared with probiotic supplementation. Colour depicts the insect species and size the sample size.

**Table 1.**
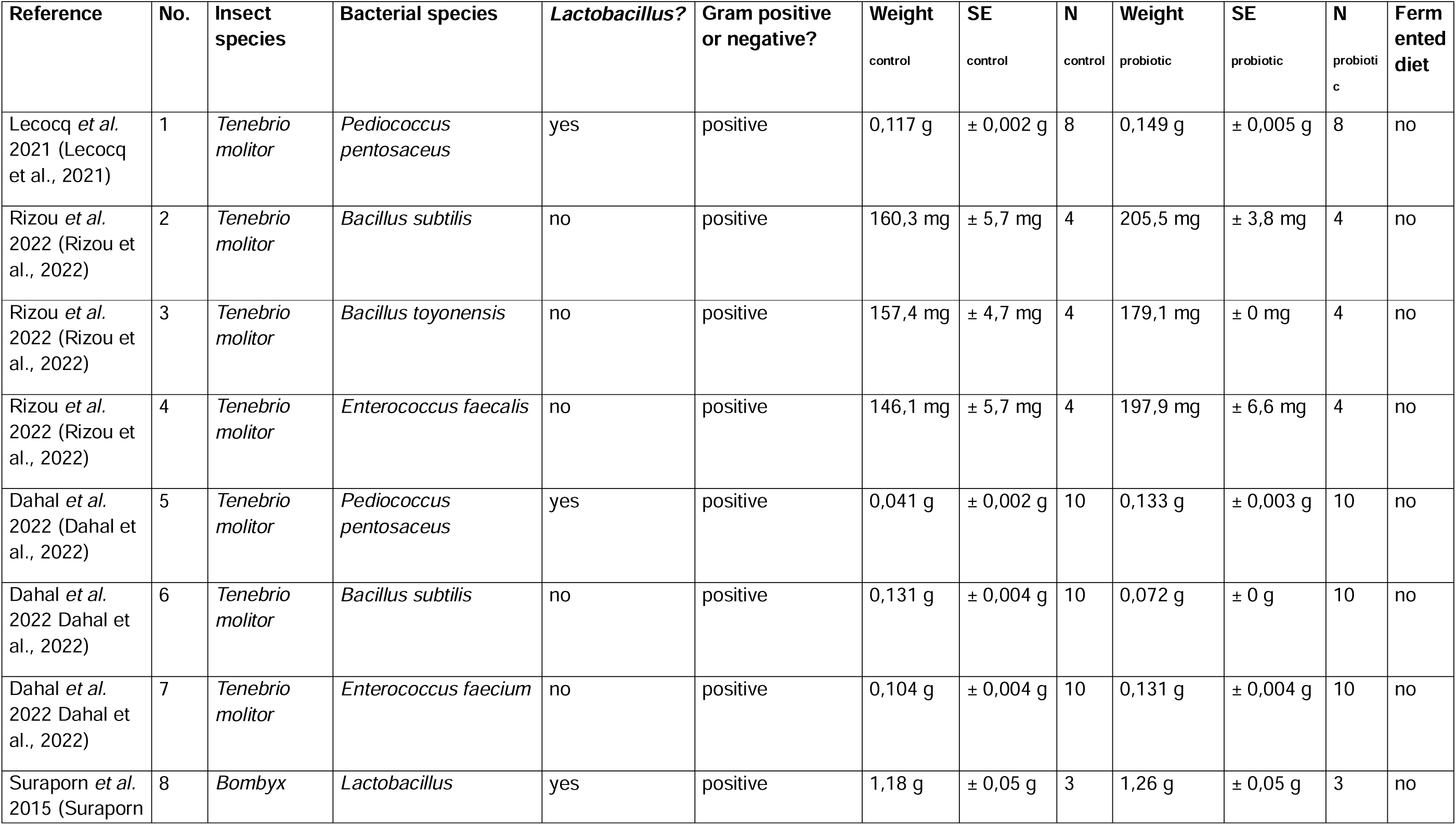
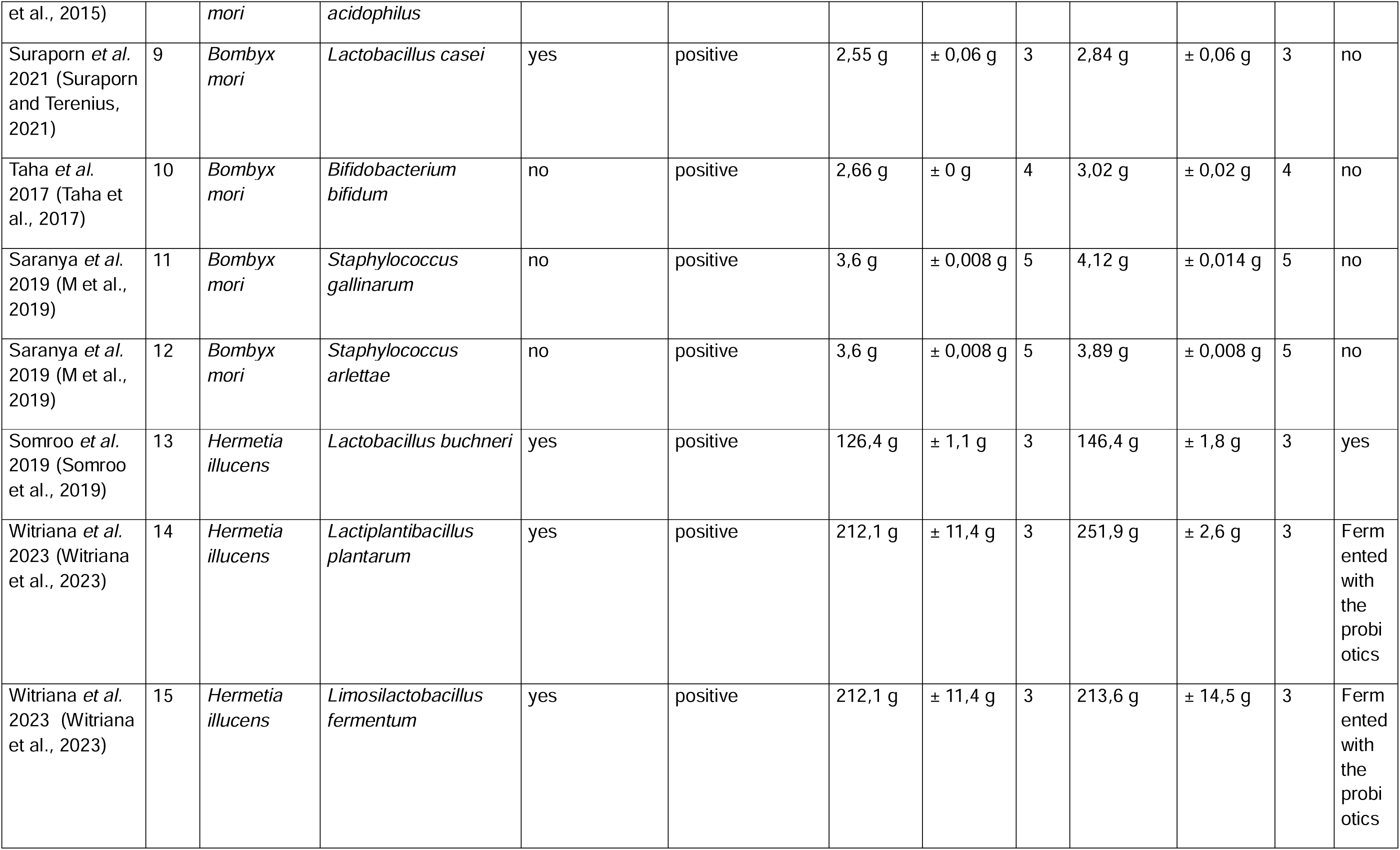
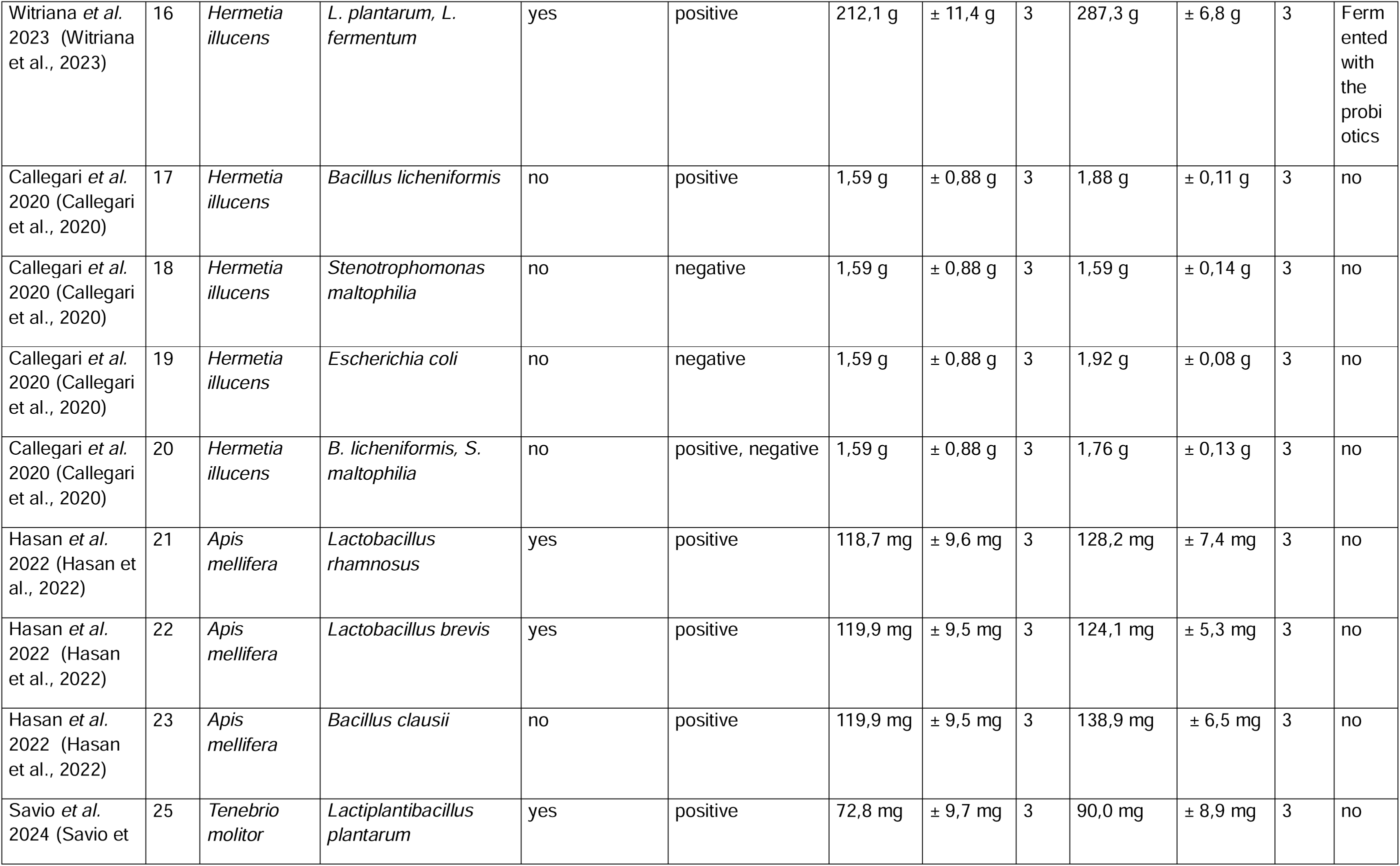
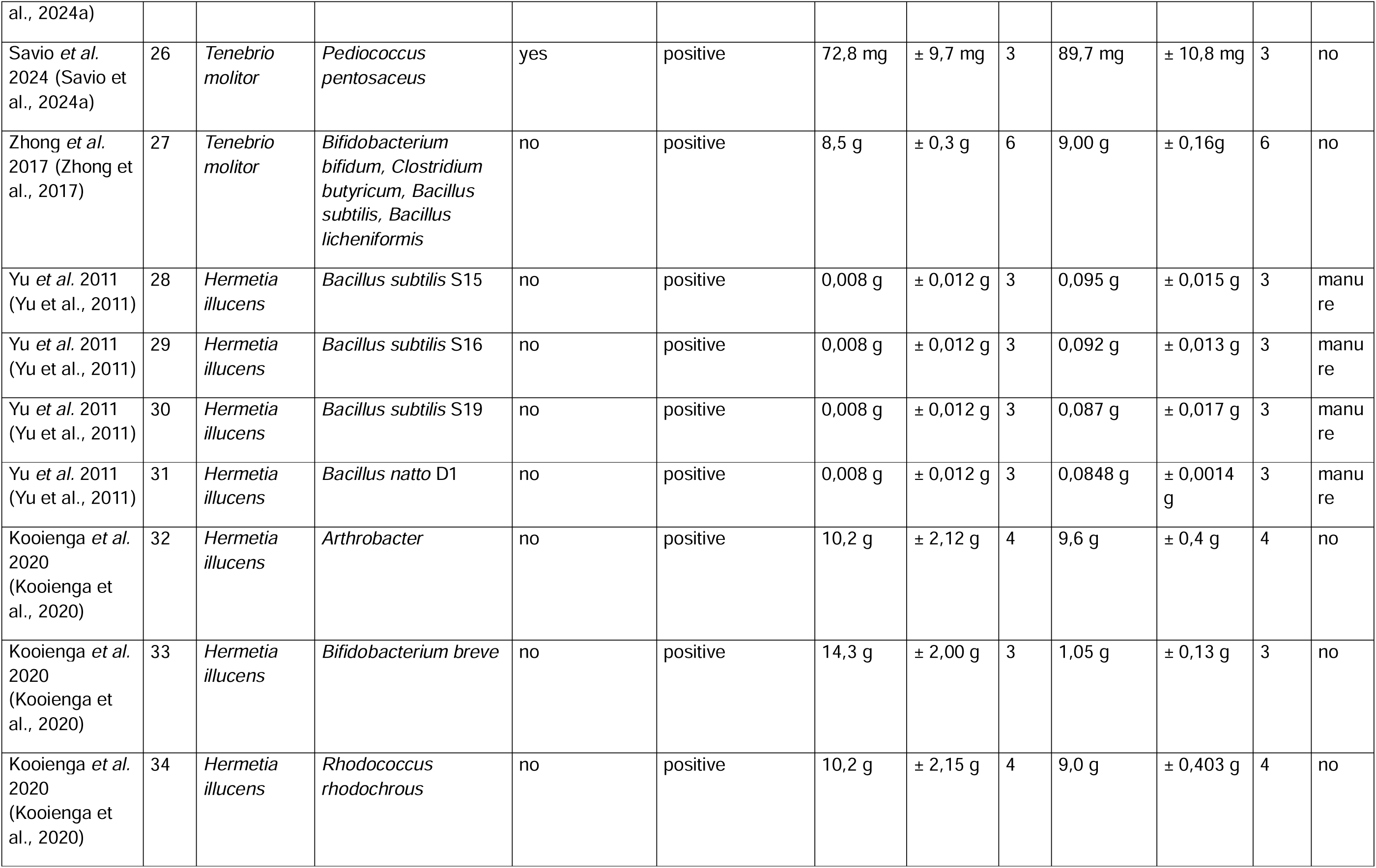
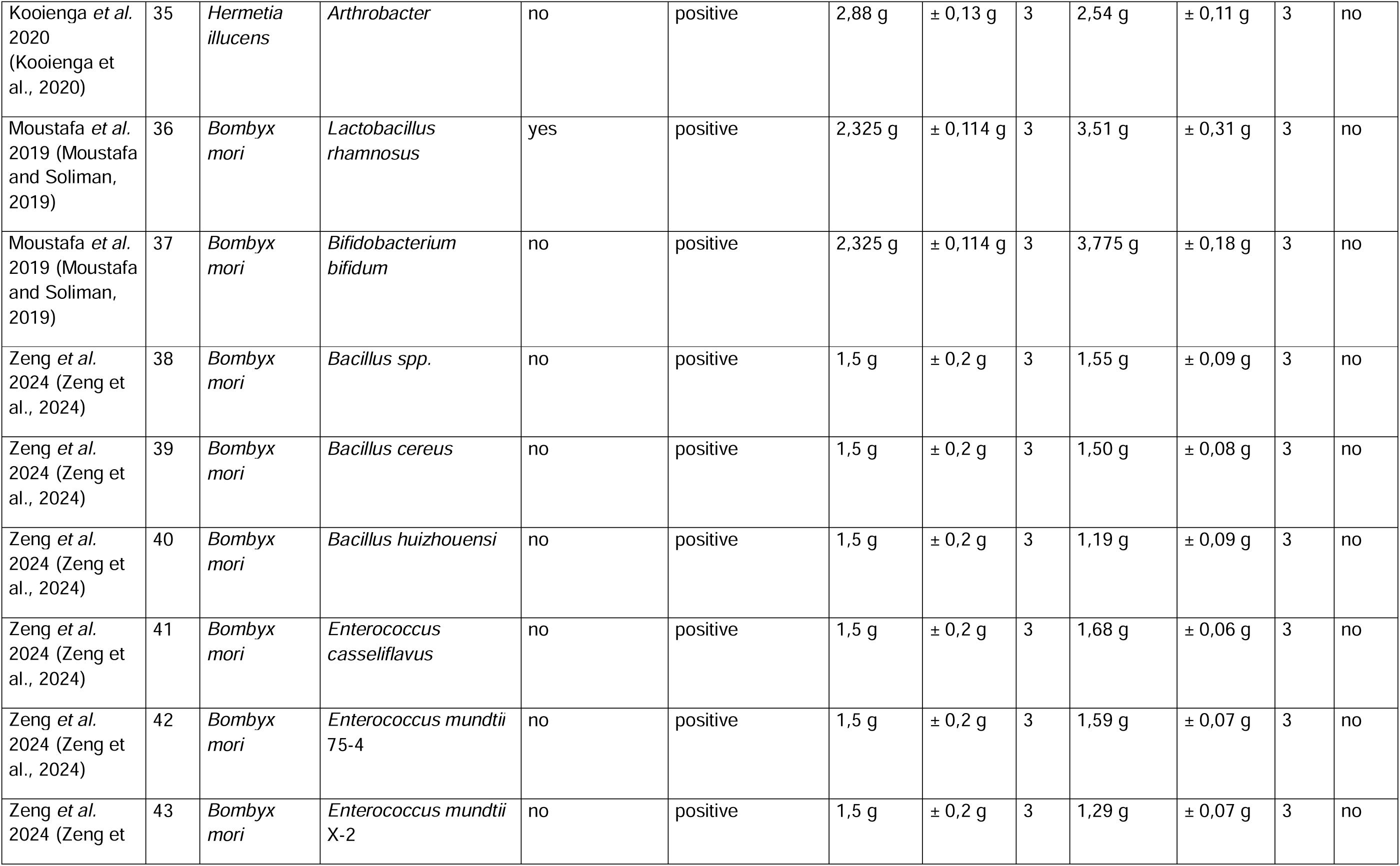
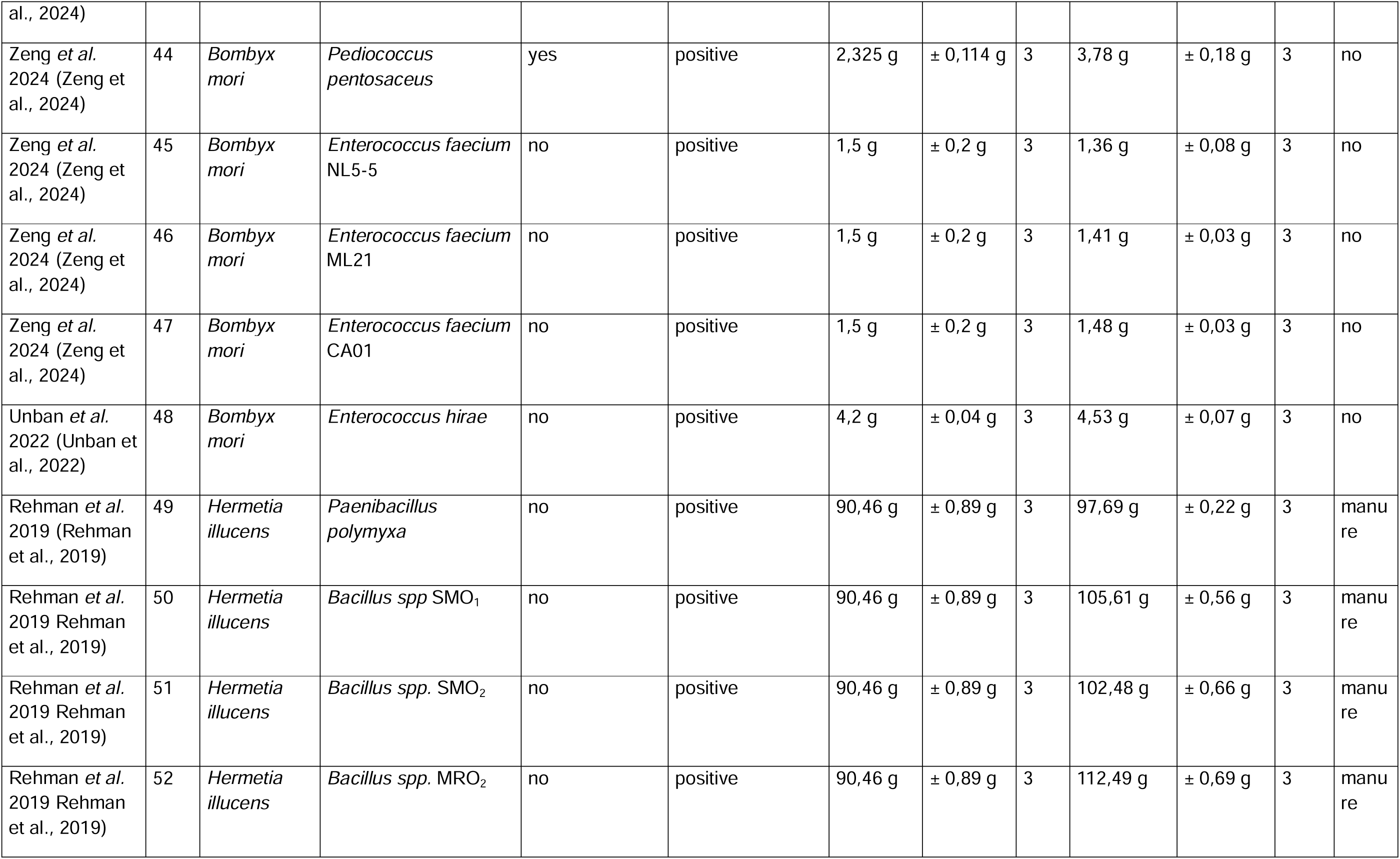
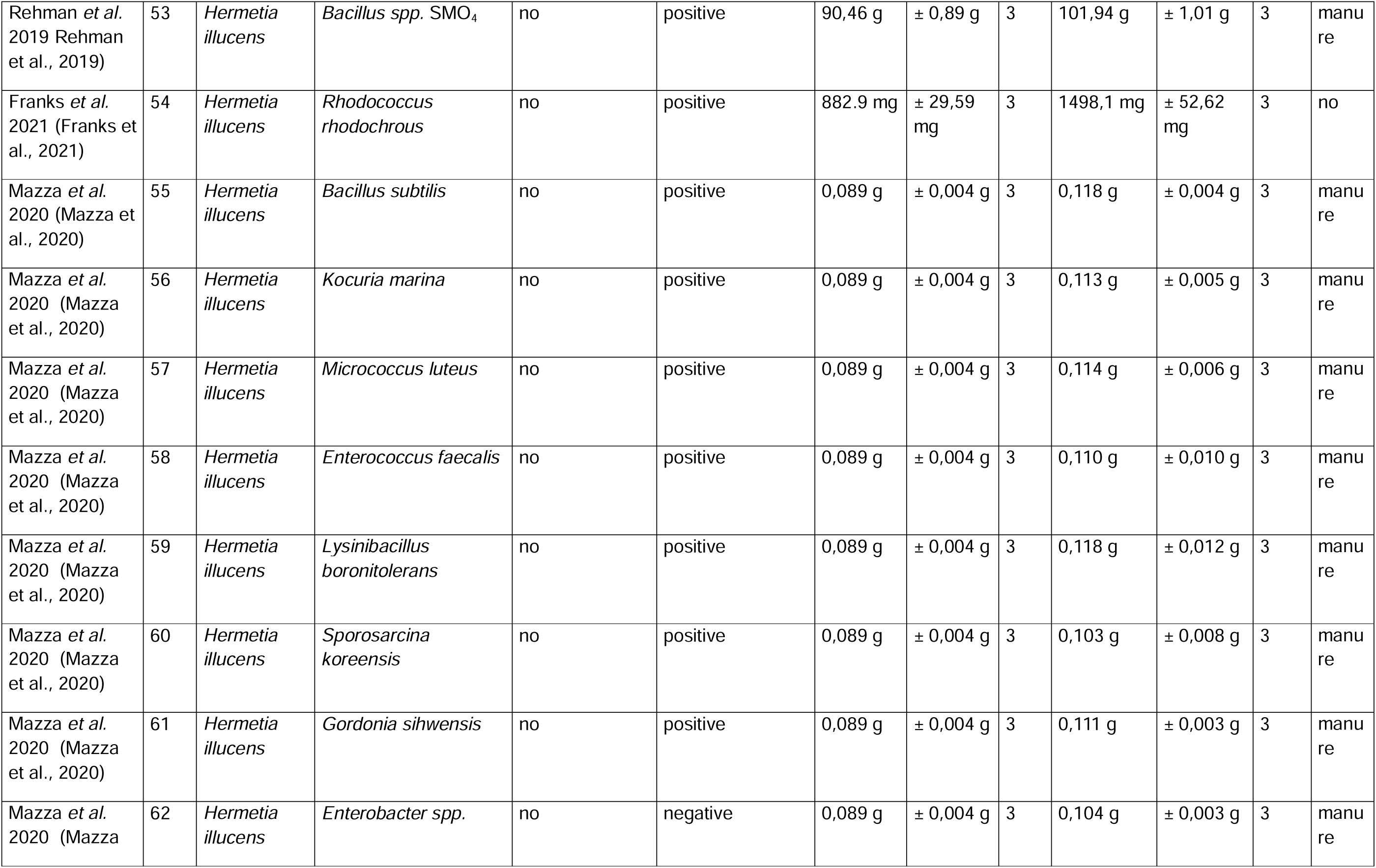
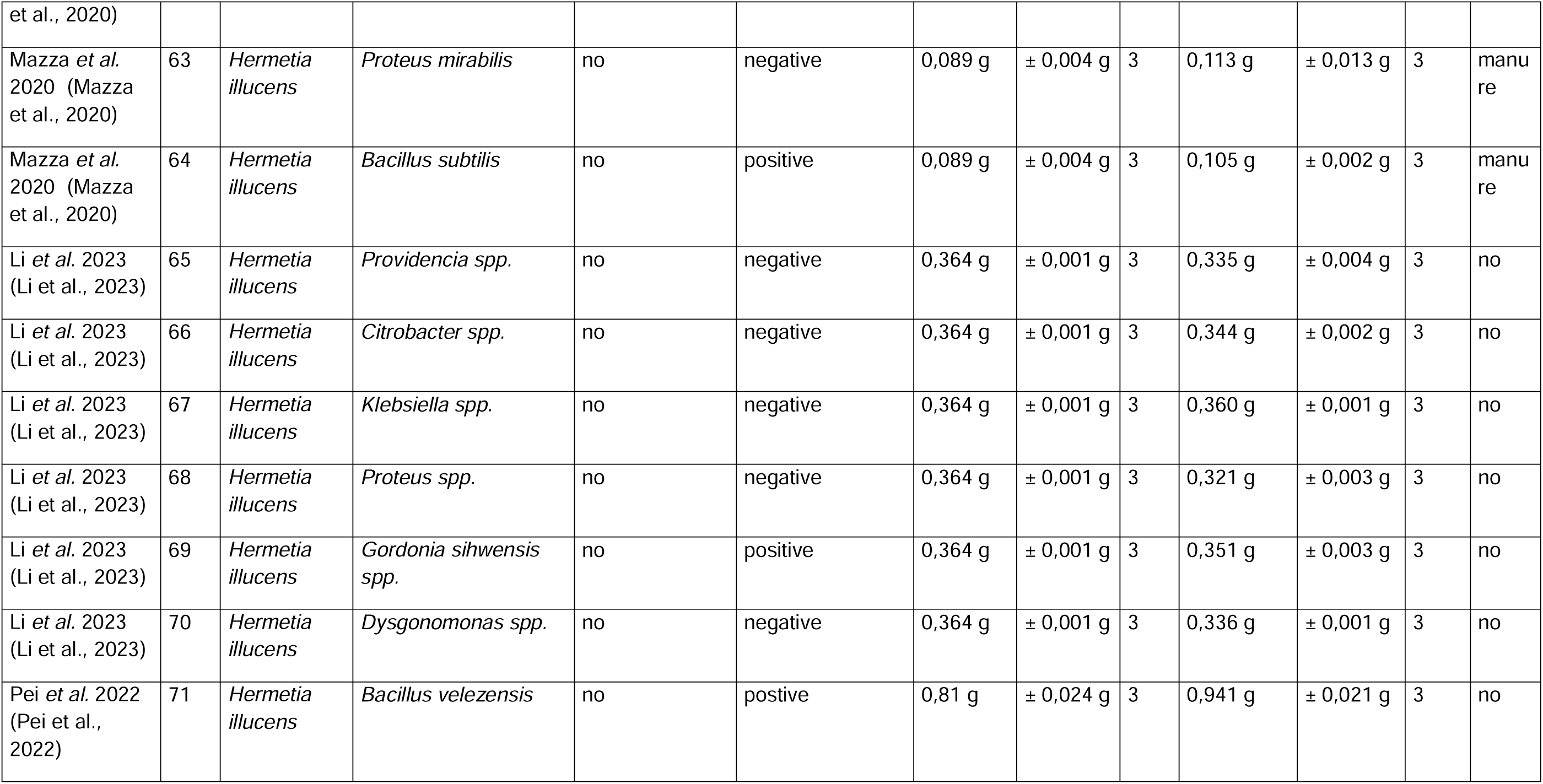
Included studies and relevant information on the insect studies included in the first meta-analysis comparing body weights.

#### Moderator variable analyses for growth rate

Insect species did not have a significant influence on the overall magnitude or direction of the effect (QM = 2.0767, d.f. = 3, p = 0.5566), however grouping studies by insect species demonstrates the disparity in numbers of studies across species (Figure 1). While we were able to include 40 datasets using *Hermetia illucens* as host, a species not yet registered in the EU as a novel food, but one of the insects bred on an industrial scale for animal feeding or agro-industrial field purposes, no studies have yet been carried out on three out of the four species that are able to be sold for food in the EU. In fact, of all four species registered as novel food in the EU, *Locusta migratoria, Acheta domesticus, Alphitobius diaperinus* and *Tenebrio molitor,* only *T. molitor* has been used a host in published studies and in just 10 of the 71 included data sets.

To test what subcategories of probiotics might be effective, we tested for differences between lactobacilli and non-lactobacilli and gram-positive and gram-negative bacteria. There was no significant difference between lactobacilli and non-lactobacilli (QM = 2.7487, d.f. = 1, p = 0.0973) or between gram-positive and gram-negative probiotics (QM =2.4363, d.f. = 2, p = 0.2958).

#### Publication bias for growth rate

To test for publication bias we first visualized the relationship between effect sizes and standard errors using a funnel plot (Figure 2A). As expected in the absence of publication bias (Field and Gillett, 2010), we saw that we tended to have larger standard errors with larger effect sizes and that we had both positive and negative effect sizes in our data set. This indicates that there was no evidence of publication bias. Furthermore, fail safe N analysis suggested that we would need to add 2393 data sets to our study to change the outcome, giving us a high level of confidence in our results.

**Figure 2.**
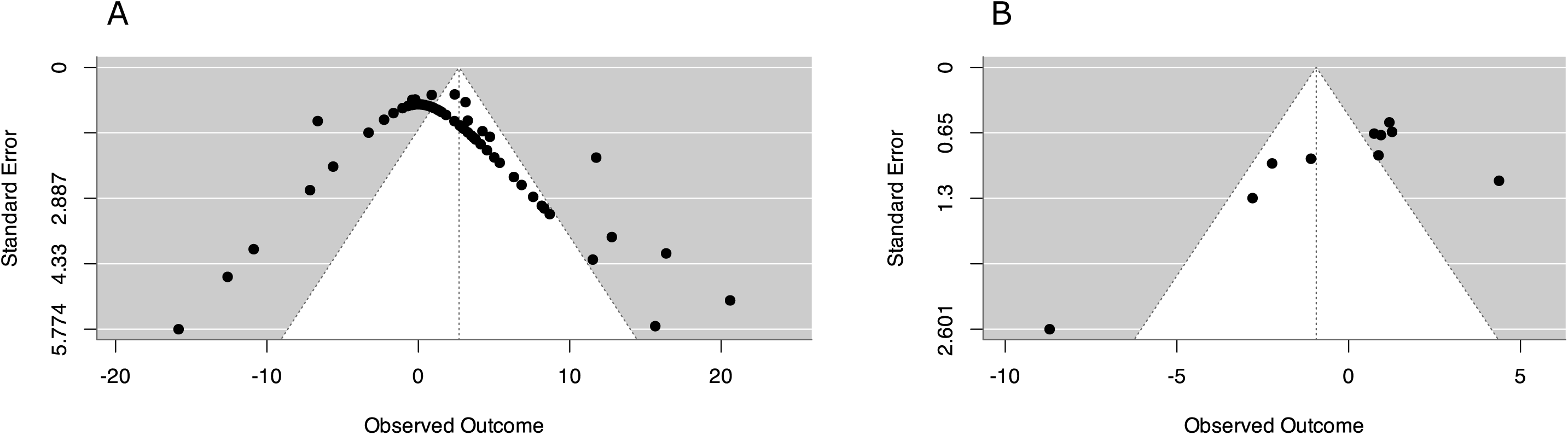
Funnel plots showing observed outcomes (effect sizes) and standard errors for: (A) the first meta-analysis comparing growth; (B) the second meta-analysis comparing microbiome diversity. The white triangle shows the expected distribution in the absence of publication bias.

### Microbiome diversity

#### Overall model for microbiome diversity

Compared to growth rate, we found fewer studies reporting data on the effects of probiotic supplementation on microbiome diversity. Nevertheless, we extracted 10 datasets from six published papers (Table 2). Overall, there was no significant effect of probiotic supplementation on microbiome diversity (estimate = -0.9449(lower ci= - 2.9918/ upper ci = 1.1020), z = -0.9048, p = 0.3656), with studies showing a range of both positive and negative effect sizes (Figure 3).

**Figure 3.**
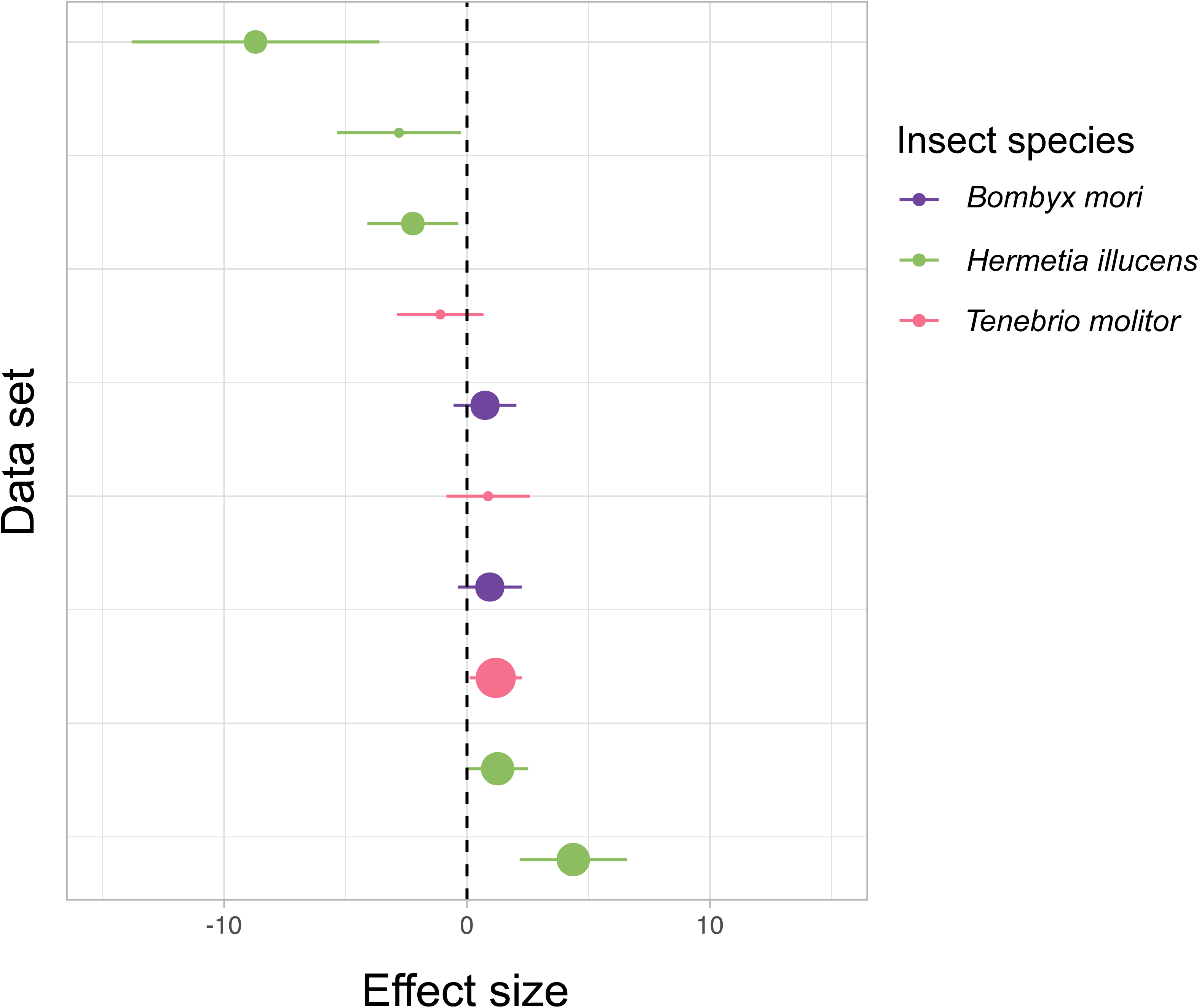
Forest plots showing effect sizes and confidence intervals for differences in insect microbiome diversity with and without probiotic supplementation with effect sizes and 95% confidence intervals for each data set included in the study, negative effect sizes show a lower microbial diversity in insects reared with probiotics than reared without and positive effect sizes show increased microbial diversity in insects reared with probiotic supplementation. Colour depicts the insect species and size the sample size.

**Table 2.**
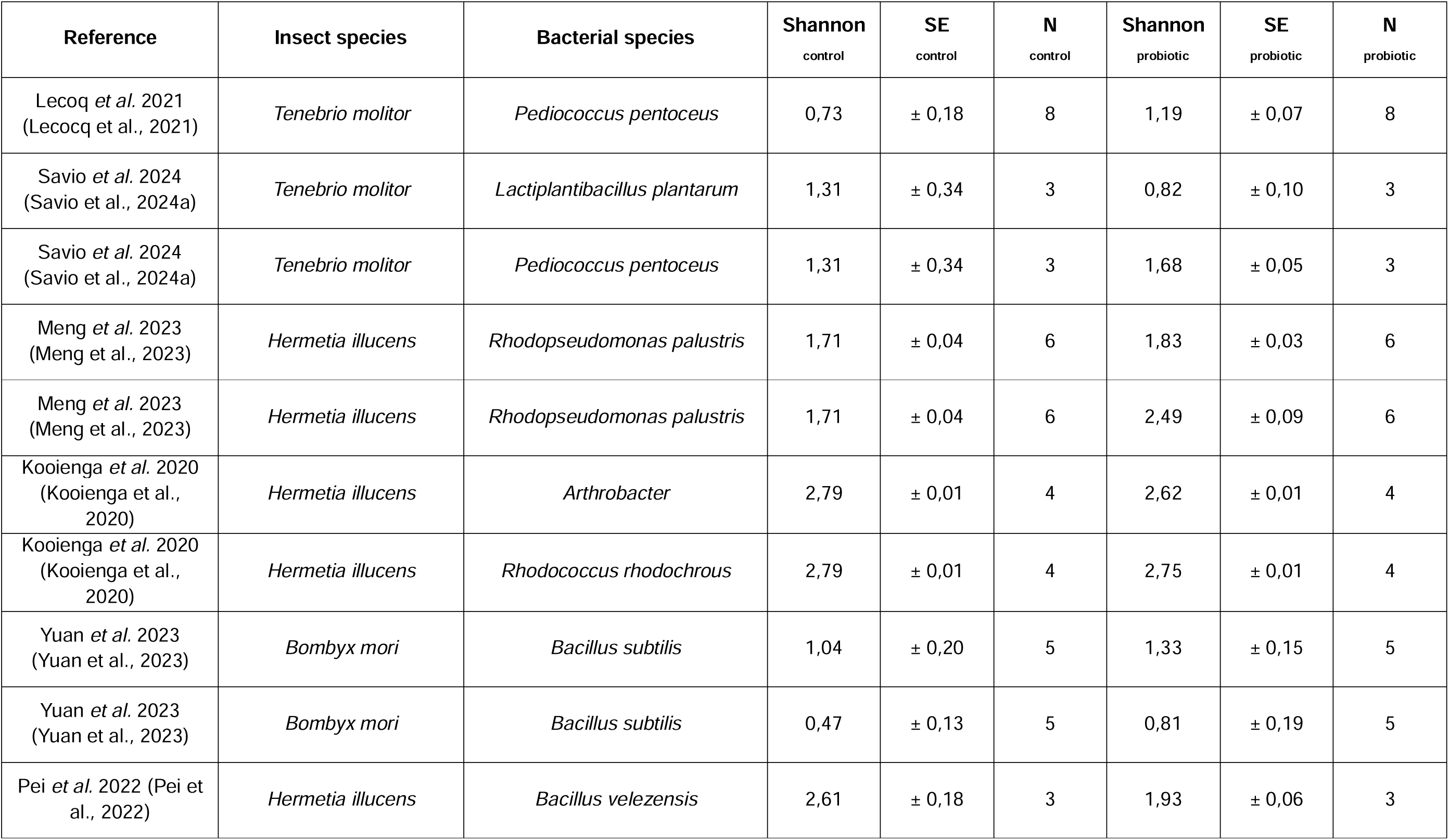
Included studies and relevant information on the insect studies included in the second meta-analysis comparing Shannon indices as an indication of bacterial community diversity.

| Reference | Insect species | Bacterial species | Shannon<br>control | SE<br>control | N<br>control | Shannon<br>probiotic | SE<br>probiotic | N<br>probiotic |
| --- | --- | --- | --- | --- | --- | --- | --- | --- |
| Lecoq *et al.* 2021<br>(Lecoq et al., 2021) | *Tenebrio molitor* | *Pediococcus pentocetus* | 0,73 | ± 0,18 | 8 | 1,19 | ± 0,07 | 8 |
| Savio *et al.* 2024<br>(Savio et al., 2024a) | *Tenebrio molitor* | *Lactiplantibacillus plantarum* | 1,31 | ± 0,34 | 3 | 0,82 | ± 0,10 | 3 |
| Savio *et al.* 2024<br>(Savio et al., 2024a) | *Tenebrio molitor* | *Pediococcus pentocetus* | 1,31 | ± 0,34 | 3 | 1,68 | ± 0,05 | 3 |
| Meng *et al.* 2023<br>(Meng et al., 2023) | *Hermetia illucens* | *Rhodopseudomonas palustris* | 1,71 | ± 0,04 | 6 | 1,83 | ± 0,03 | 6 |
| Meng *et al.* 2023<br>(Meng et al., 2023) | *Hermetia illucens* | *Rhodopseudomonas palustris* | 1,71 | ± 0,04 | 6 | 2,49 | ± 0,09 | 6 |
| Kooienga *et al.* 2020<br>(Kooienga et al., 2020) | *Hermetia illucens* | *Arthrobacter* | 2,79 | ± 0,01 | 4 | 2,62 | ± 0,01 | 4 |
| Kooienga *et al.* 2020<br>(Kooienga et al., 2020) | *Hermetia illucens* | *Rhodococcus rhodochrous* | 2,79 | ± 0,01 | 4 | 2,75 | ± 0,01 | 4 |
| Yuan *et al.* 2023<br>(Yuan et al., 2023) | *Bombyx mori* | *Bacillus subtilis* | 1,04 | ± 0,20 | 5 | 1,33 | ± 0,15 | 5 |
| Yuan *et al.* 2023<br>(Yuan et al., 2023) | *Bombyx mori* | *Bacillus subtilis* | 0,47 | ± 0,13 | 5 | 0,81 | ± 0,19 | 5 |
| Pei *et al.* 2022 (Pei et al., 2022) | *Hermetia illucens* | *Bacillus velezensis* | 2,61 | ± 0,18 | 3 | 1,93 | ± 0,06 | 3 |

#### Moderator variable analysis for microbiome diversity

Again here, insect species did not have an impact on the magnitude or direction of the effect (QM = 1.0816, d.f. = 2, p =0.5823), although the breakdown also highlighted that in diversity measures, just as in growth rate, there is a bias in the insect species in which research has been carried out, with the majority of the research having been done on the black soldier fly, *H. illucens (*Figure 3). For this analysis there were an insufficient number of studies to meaningfully test further moderator variables.

#### Publication bias for microbiome diversity

We plotted a funnel plot to visualize potential publication bias for microbiome diversity, however, due to the small number of datasets meeting the criteria for inclusion in the meta-analysis for microbiome diversity it is difficult to draw clear conclusions (Figure 2B).

#### Inhibitory effects of probiotics against pathogen infection

For the benefits of probiotics to be maximised, we would ideally supplement farmed insects with probiotics that not only stimulate growth and gut health but that also provide protection against pathogens. Unfortunately, to date there are too few studies to carry out a meaningful meta-analysis on this specific question. To our knowledge published work on the defensive properties of probiotics against pathogens has been mainly carried out in *T. molitor* against entomopathogens. The few published studies present mixed, yet potentially promising results. Lecocq et al (Lecocq et al., 2021) showed that *Pediococcus pentoceus* has inhibitory effects in vitro against a range of potentially relevant entomopathogens. Building on this, Dahal et al (Dahal et al., 2022) showed in vivo that supplementation of *T. molitor* with *P. pentoceus*, not only enhances growth but also seems to provide some degree of protection to *T. molitor* against mortality induced by the highly virulent entomopathogenic fungus *Metarhizium brunneum*, although the potential probiotic *B. subtilus* trends towards negatively impacting survival upon infection (Dahal et al., 2022). The positive results with *P. pentoceus* were, however, further supported by Savio et al (Savio et al., 2024a) who showed that *P. pentoceus* seems to provide protection to *T. molitor* against coinfection with *Bacillus thuringiensis* and *Metarhizium brunneum*, although there are some differences in the outcomes of the studies, which suggests a degree of context dependency.

There is also evidence of probiotic protection against pathogens in the silk worm *Bombyx mori*. Here supplementation with *Lactobacillus casei* resulted in protection against infection by the microsporidian *Nosema bombycis* (Suraporn and Terenius, 2021) and supplementation with *Lactobacillus lactis* protection against infection with *Pseudomonas aeruginosa* (Nishida et al., 2016). The mechanism of *Lactobacillus* induced protection is thought to be via bacterial induced activation of the immune system (Nishida et al., 2016).

## Discussion

There is a rapidly growing literature base on the supplementation of insects with probiotics. Despite this growing literature, however, we only were able to extract a sufficient number of data sets for meta-analysis of two measures: insect growth and microbiome diversity. While probiotic supplementation significantly enhances insect growth across tested species, probiotics do not have a clear effect on microbial diversity, with studies showing both positive and negative effects. Furthermore, our data clearly highlight some substantial literature gaps where more data are urgently needed. Most striking is the restricted number of insects in which these kinds of test have been carried out and the lack of studies on three of the four insect species registered as novel food in the EU.

Across the data that are available we saw a very strong and robust positive effect of probiotic supplementation on insect growth. Rearing insects on probiotics resulted in heavier insects. By decomposing indigestible fibres, producing essential nutrients, and enabling metabolic and signalling processes (Chabanol and Gendrin, 2024), the probiotics play a key role in the nutrition and development of the insect host.

Moreover, they may aid the insects’ ability to cope with stress factors (temperature, toxins, chemicals, pathogens, etc.). This is a phenomenon known from other mutualisms between insects and microbes (Armitage et al., 2022; Ford and King, 2016; Vorburger and Perlman, 2018).

We did not see a significant effect of probiotic supplementation on microbiome diversity albeit based on a very small sample size. Although microbiota diversity, often in the form of a Shannon index, is a commonly reported measure in probiotic supplementation studies, it is difficult to judge what exactly a reduced or increased Shannon index means for insect health (Johnson and Burnet, 2016). It is often assumed that high diversity means a healthy microbiome, likely due do associations with low diversity and human disease (Huttenhower et al., 2012). However, when supplementing with probiotics at high doses we may, to an extent, replace the unknown microbiota, which may be healthy or unhealthy, with bacterial species known to benefit health. Thus, reduced diversity may be due to replacement with the probiotic and not necessarily indicate poor or even reduced health.

The increase in growth observed across studies when insects are supplemented with probiotics supports the idea that probiotics enhance insect health. What is more this benefit to growth results in increased yields, bringing additional economic advantages. In spite of the indication of enhanced health coming from the growth rate data, however, we found that very little data were available on the susceptibility to pathogens of probiotic supplemented insects. This is in spite of the fact that probiotics are known to have protective effects in aquaculture settings (Chauhan and Singh, 2019; Kuebutornye et al., 2020; Sharifuzzaman and Austin, 2017), and are even being considered as an alternative to antibiotics in livestock farming (Leistikow et al., 2022). Furthermore, the use of probiotics or “defensive microbes” has been advocated as a method of pathogen control in applied settings in general (Ford and King, 2016) as well as for the specific case of edible insects (Grau et al., 2017; Savio et al., 2022). The studies that have been carried out looking at potential protection of insects from pathogen attack by probiotics do show promising results, with evidence that some probiotics inhibit the growth of pathogens (Dahal et al., 2022; Lecocq et al., 2021; Nishida et al., 2016; Savio et al., 2024a; Suraporn and Terenius, 2021). Much more data are needed, however, to fully explore the potential protective effects of candidate probiotics against pathogen infection across the full range of edible insect species.

Harnessing probiotics as an alternative to antibiotic treatment to prevent the establishment and spread of disease within insect cultures presents a strategy through which we may avoid the overuse of antibiotics seen in livestock rearing. However, this strategy is not without risks. We know that probiotics can also be a source of AMR genes (Daniali et al., 2020; Radovanovic et al., 2023; Savio et al., 2022; Tóth et al., 2021), which could mean that if probiotic treatment is unsuccessful and antibiotics do have to be used, efficacy is impacted and the presence of probiotics harbouring such genes may even enhance the rate of AMR evolution. On the other hand, we can use this knowledge to ensure that probiotics are bred exclusively from AMR free strains. Also here, this is a question that urgently needs to be explored both experimentally and theoretically as the scale of insect production for food and feed increases.

The microbiome is critically associated with a range of behavioural and physiological function and how the immune response might be linked to probiotic supplementation remains unclear. There has been much discussion as to how the host immune system balances the need to allow the colonisation of mutualistic micro-organisms while simultaneously fighting off pathogens (Betts et al., 2016; Hanson, 2024), as research in the field develops, it is important we consider how potential immune effects of microbiome manipulation may impact mass rearing.

Finally, it is important to consider the microbial composition of the feeding material given to insects grown as food and feed. Our literature research showed that particularly black soldier fly larvae are being fed a range of microbe rich (Mazza et al., 2020; Rehman et al., 2019) and fermented material (Somroo et al., 2019; Witriana et al., 2023). Fermentation can result in enrichment of probiotics (Heller, 2001). Given that fermentation of vegetable and other food waste provides an opportunity to utilise organic side streams and increase the climate friendliness of insects as food and feed further, the impact of these foods on insect health and growth warrants further investigation (Antunes et al, unpublished data).

Our systematic analysis shows that a strong body of research is developing on the topic of probiotic supplementation in edible insects. As research continues, we highlight six key questions that remain to be addressed. These are: 1) Do growth enhancing probiotics also provide protection against pathogens? 2) Does probiotic supplementation reduce or increase the risk of AMR? 3) Do the effects we see in the insect species studied so far extent to other edible insect species, particularly those being sold as novel food in the EU? 4) Does probiotic supplementation impact insect behaviour under mass rearing conditions? 5) How does probiotic supplementation affect the insect immune system? 6) What role do diet substrates, and their processing techniques play in shaping probiotic diversity and stability within the insect gut? Our hope is that continued research in this field, addressing the highlighted questions has the potential to greatly improve the sustainability and efficiency of insect rearing for food and feed.

## Conclusions

We have provided the first systematic review of the quantitative impacts of probiotic supplementation on edible insects. Overall probiotics tend to boost insect growth, which suggests that insect health is enhanced through supplementation. However, there is no clear effect on microbiome diversity. There are also two clear literature gaps highlighted by our study. First of all, the taxonomic restriction on the insect species that have been studied to date. If probiotic supplementation is to be considered as an implementation strategy to support insect health in commercial rearing, it is important that those insects with the highest commercial potential are testing. Secondly, our study highlights a lack of studies investigating the protective effects of probiotics against pathogenic infection. As the focus on probiotics in the edible insect industry grows, we hope these knowledge gaps will be filled.

